# Evolving resistance patterns in *Tetranychus urticae* and *Bemisia tabaci* in Greece

**DOI:** 10.1101/2025.05.31.656940

**Authors:** Anastasia Kampouraki, Aris Ilias, Kiriaki-Maria Papapostolou, Stella Malliaraki, Ioannis Pirgiannakis, Evangelia Karakosta, John Vontas, Anastasia Tsagkarakou, Konstantinos Mavridis

## Abstract

Pesticide resistance in agricultural pests poses a significant challenge to sustainable crop protection. In this study, we assessed the current resistance status of two key pests, Tetranychus urticae and Bemisia tabaci, in major agricultural regions of Greece. A total of 25 field populations were collected between 2023 and 2025 and tested using single-dose toxicity bioassays and droplet digital PCR (ddPCR) for known resistance-associated target-site mutations. The results revealed significant variability in susceptibility to multiple acaricides and insecticides, with several T. urticae populations displaying reduced sensitivity to abamectin, hexythiazox, and fenpyroximate. Resistance mutations such as I321T (GluCl3), I1017F (CHS1), F1538I (VGSC), and G126S (cytb) were detected at high frequencies in multiple populations, indicating multi-resistant phenotypes. In B. tabaci, widespread occurrence of F331W (ace1), L925I/T929V (VGSC), and A2083V (acc) mutations confirmed evolving resistance to organophosphates, pyrethroids, and ketoenols, respectively. Our findings underscore the ongoing evolution of resistance in these pests and highlight the need for integrated management strategies that include regular resistance monitoring and judicious pesticide use. This study provides data to guide evidence-based pest control and supports the implementation of sustainable Integrated Pest Management (IPM) practices in Greece.

## 1. Introduction

The emergence and fast spread of pesticide resistance in agricultural pests is a major threat to crop protection and food production. Resistance has been recorded in many arthropod species around the world and is often linked to the repeated use of insecticides. Two of the most important pests globally, the two-spotted spider mite (*Tetranychus urticae*) and the silverleaf whitefly (*Bemisia tabaci*), are especially known for their ability to develop resistance to several groups of pesticides. According to the Arthropod Pesticide Resistance Database (Mota-Sanchez, D. and J.C. Wise, 2025), there are currently 558 reported cases of resistance for *T. urticae* and 898 for *B. tabaci*.

In Greece, these pests are serious threats for many vegetable, fruit, and ornamental crops, both in greenhouses and in open fields. Integrated Pest Management (IPM) strategies are increasingly promoted as a holistic approach to pest control, encouraging the use of biological control agents, crop rotation, resistant cultivars, and environmentally friendly alternatives. Among these, the use of “green chemicals” or biorational pesticides—products derived from natural sources such as plant extracts, microbial metabolites, or naturally occurring fatty acids—has gained ground in recent years. These compounds are considered to have low toxicity to non-target organisms and a lower risk for resistance development, and their integration into IPM programs is actively encouraged (Copping and Menn, 2000; Isman, 2006; Paspati et al., 2023). Nevertheless, chemical control remains the main method currently used in the field, particularly against *T. urticae* and *B. tabaci*.

Several pesticide groups with different Modes of Action (MoA) are applied, including neonicotinoids, sulfoximines, and butenolides (which act on nicotinic acetylcholine receptors), abamectin (targeting the glutamate-gated chloride channels GluCls), pyriproxyfen (a juvenile hormone analog), hexythiazox (a mite growth inhibitor MGI targeting the chitin synthase CHS1), Mitochondrial Electron Transport Inhibitors (METI) like pyridaben, fenpyroximate, and acequinocyl, and diamides (affecting the ryanodine receptor). Due to their frequent use, resistance has developed in many field populations of *T. urticae* and *B. tabaci* (Dermauw et al., 2012; Ilias et al., 2012, 2015; Khajehali et al., 2010; Mavridis et al., 2022a, 2022b; Papapostolou et al., 2021; Riga et al., 2014, 2015; Roditakis et al., 2005, 2006, 2011; A. Tsagkarakou et al., 2009). One of the main mechanisms of resistance in these pests is the presence of mutations in the target sites of pesticides.

In *T. urticae*, several known mutations have been associated with resistance: I1017F in the CHS1 gene (for MGIs like hexythiazox and etoxazole), F1538I and L1024V in the voltage-gated sodium channel (VGSC) gene (linked with pyrethroid resistance), G126S, H92R, P262T and S141F in the mitochondrial cytochrome b gene (for resistance to METI-I and METI-III acaricides), and G314D, G326E, and I321T in the GluCl genes (for abamectin resistance) (De Rouck et al., 2023; Dermauw et al., 2012; Riga et al., 2017; Van Leeuwen et al., 2010, 2008).

In *B. tabaci*, several resistance mutations have also been reported. These include the F331W mutation in the ace1 gene (linked to resistance to organophosphates and carbamates) (Alon et al., 2008), the L925I and T929V mutations in the VGSC gene (for pyrethroid resistance) (Morin et al., 2002; Roditakis et al., 2006), and the A2083V mutation in the acc gene (associated with resistance to ketoenols like spiromesifen) (Lueke et al., 2020). In addition, the A387G mutation in the cytochrome P450 gene CYP6CM1 has been associated with neonicotinoid resistance (Pym et al., 2023).

To improve monitoring and resistance detection, droplet digital PCR (ddPCR) was recently developed as a highly sensitive molecular method to monitor the presence and frequency of resistance mutations with high accuracy and sensitivity (Mavridis et al., 2022a, 2022b), in pooled insect and mite DNA samples.

In this study, we present updated resistance data for *T. urticae* and *B. tabaci* field populations collected from major agricultural regions of Greece. Using a combination of single-dose toxicity bioassays and ddPCR diagnostics, we monitored the current resistance status of these key pests in Greece, aiming to guide efficient and sustainable pest management.

## 2. Materials and Methods

### 2.1 Tetranychus urticae and Bemisia tabaci field samples

A total of 19 field populations (Table 1) of *Tetranychus urticae* (Tu1–Tu19) and 6 populations of *Bemisia tabaci* (Bt1–Bt6) were collected from different areas of Greece between September 2023 and January 2025. Sampling was carried out from infested leaves of various host plants (vegetables, citrus, ornamentals), collected from greenhouse and open-field conditions. Collection areas included major agricultural zones such as the Peloponnese (Achaea, Argolida, Messinia) and Crete (Heraklion and Lasithi prefectures).

**Table 1.**
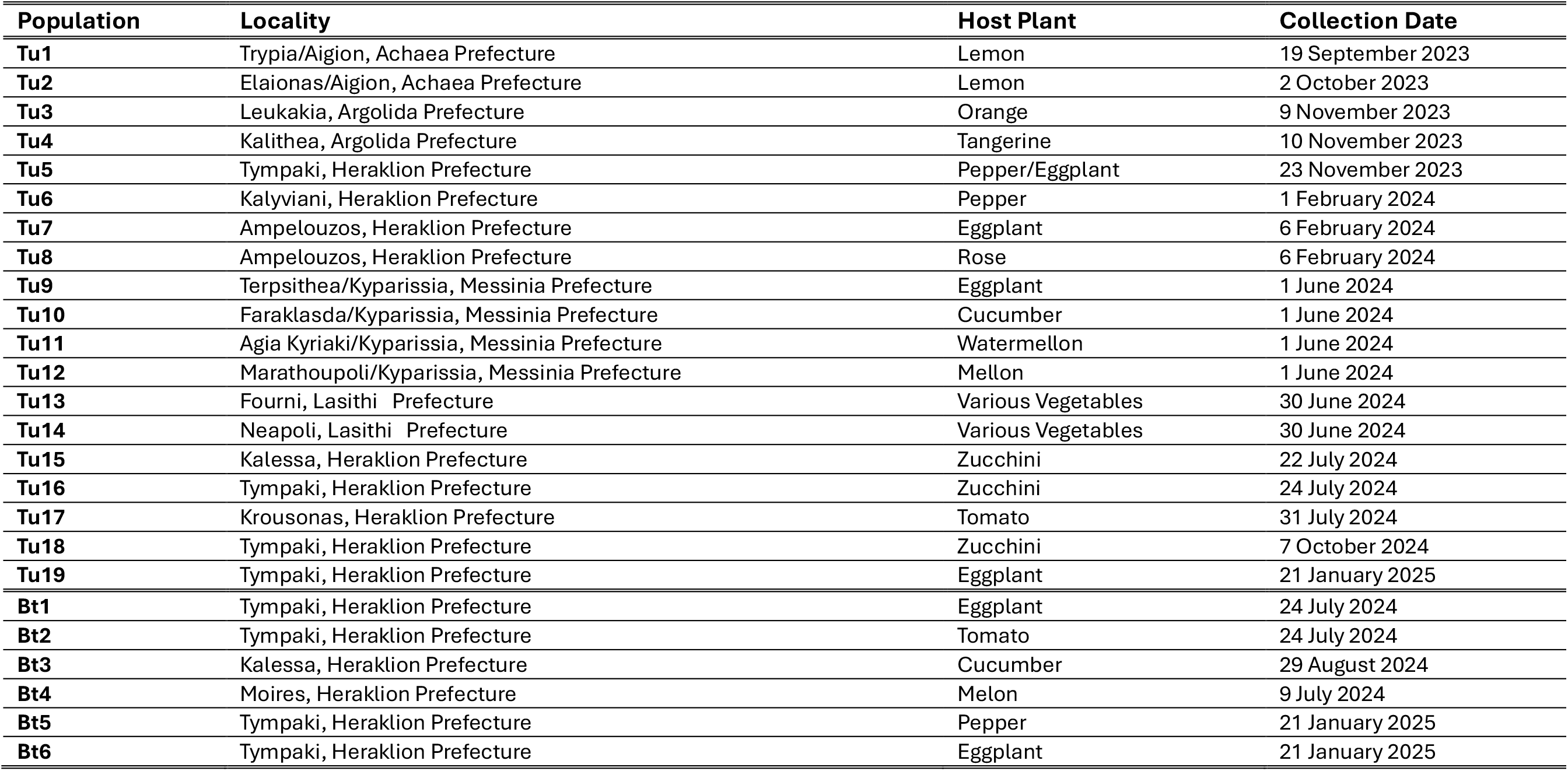
Characteristics of the *T. urticae* field populations collected from Crete.

The *T. urticae* populations were collected from citrus crops (lemon, orange, tangerine), vegetables (cucumber, zucchini, tomato, eggplant, melon, watermelon, pepper), and ornamental crops (rose). For several populations, pesticide application history was known and included a wide range of compounds. In particular, Tu5 (from pepper/eggplant in Tympaki) had a history of treatments with spinetoram, spinosad, metaflumizone, formetanate, and acrinathrin. Tu9 (from eggplant in Messinia) was treated with pyriproxyfen, acetamiprid and flonicamid, while Tu11 (from watermelon in the same area) had been treated with abamectin. Tu12 (from melon) was exposed to acetamiprid and chlorantraniliprole, Tu16 (zucchini) to pyriproxyfen, and Tu18 (zucchini) to hexythiazox and acequinocyl and fatty acid potassium salts. For the rest of the populations, either no chemical treatments had been applied (Tu1, Tu4, Tu6, Tu7, Tu8, Tu17, Tu19) or the pesticide history was unknown (Tu2, Tu3, Tu10, Tu13, Tu14, Tu15).

The *B. tabaci* populations were collected mainly from solanaceous and cucurbit crops such as eggplant, tomato, cucumber, melon, and pepper. In this case, pesticide use was recorded for several samples. Bt1 (eggplant, Tympaki) had been exposed to flonicamid, acetamiprid, pyridaben, pyriproxyfen, and sulfoxaflor; Bt2 (tomato, Tympaki) to azadirachtin, acetamiprid, and pyriproxyfen; Bt5 to cyantraniliprole, sulfoxaflor, pyriproxyfen, pyridaben and deltamethrin. Although the Bt6 (eggplant, Tympaki) field was known to be untreated during the sampling period, it had a heavy history of chemical applications. In the remaining cases (Bt3, Bt4), the treatment history was unknown.

### 2.2 Toxicity assays

Four formulated acaricides/insecticides of different MoA, abamectin (Vertimec 18EC), bifenazate (Floramite 240SC), fenpyroximate (Kento 5.12SC) and hexythiazox (Nissorun 25SC), and three alternative “green” products, Requiem, Eradicoat, and Flipper were used for toxicity assays, using diagnostic recommended field doses. Adulticidal bioassays were conducted with 25 2-3 days old adult female mites which were transferred on the upper side of 9 cm^2^ square-cut kidney bean leaf discs on wet cotton wool. The plates were sprayed with 1 ml of spray fluid at 1 bar pressure with a Potter Spray Tower (Burkard Scientific, UK) to obtain a homogenous spray film. The toxic effects of hexythiazox were assessed with egg bioassays, by having 20 adult females laying eggs on a leaf disc for 5h, as previously described. After spray treatment, leaf discs were placed at 25 ± 0.5 °C, 60% RH and 16:8 h (light:dark) photoperiod. The recommended field dose (RD) for each acaricide was applied in toxicity assays. Three replicates were sprayed per dose and, as a control, deionized water was used. Mortality was assessed after 72h for abamectin, bifenazate, fenpyroximate, Requiem, Eradicoat and Flipper. Adult mites were considered alive if they could walk twice the distance of their body size after being prodded with a fine hairbrush. Mites treated with hexythiazox were considered unaffected, if they displayed the same developmental stage as the water treated control at the time of scoring. The percentage mortality resulting from exposure to each insecticide was calculated after being corrected for control mortality. The percentage mortality for each insecticide was corrected using Schneider-Orelli’s formula, i.e., Corrected % mortality = [(mortality % in treatment-mortality % in control) / 100 − mortality % in control)] × 100.

### 2.3 Genomic DNA isolation and quantification

Genomic DNA extractions were performed on pools of adult mites, following the CTAB method as previously described (Navajas et al., 1999). The concentrations of extracted gDNAs were determined with the Qubit™ dsDNA BR Assay (Invitrogen, Carlsbad, CA) on a Qubit fluorometer 2.0 (Invitrogen, Carlsbad, CA), in order to obtain measurements specific for double stranded DNA (dsDNA). The mean ± SE dsDNA concentration for *T. urticae* pools was 12.2 ± 3.76 (range: 4.6–25.0 ng μL–1), while for the B. *tabaci* pools 77.9 ± 19.85 (range: 11.0–140.0 ng μL–1).

### 2.4 Droplet Digital PCR (ddPCR) Assays

Droplet digital PCR (ddPCR) was performed using the QX200 Droplet Digital PCR System (Bio-Rad, Hercules, CA, USA), as previously described (Mavridis et al., 2022b, 2022a) Each 20 μL reaction contained 1× ddPCR Supermix for Probes (no dUTP) (Bio-Rad), 5 U of EcoRI-HF® restriction enzyme (New England Biolabs), 5 ng of genomic DNA, and primers and probes specific to each mutation. Probes were labeled with FAM for mutant alleles and HEX for wild-type alleles.

Droplets were generated with the QX200 droplet generator and thermocycled under the following conditions: 95 °C for 10 min, followed by 50 cycles of 94 °C for 30 s and 54–60 °C for 1 min, and a final step at 98 °C for 10 min. After amplification, droplets were read on the QX200 droplet reader, and data were analyzed using QuantaSoft™ Analysis Pro Software (v1.0.596). The mutant allelic frequency (MAF) was calculated as: %MAF = [Mutant copies / (Mutant + Wild-type copies)] × 100.

The limit of detection (LoD) was determined using synthetic gBlock™ controls and reached 0.1–0.2%, depending on the assay. This sensitivity allows for reliable detection of low-frequency resistance alleles in pooled samples (Mavridis et al., 2022a, 2022b).

### 2.5 Statistical analyses

The limit of detection (LoD) for each ddPCR assay was defined as the lowest MAF that could be reliably detected with accuracy calculated according to (Lavebratt et al., 2004).

## 3. Results

### 3.1 Toxicity assays

Toxicity assays were conducted on twelve field populations of *Tetranychus urticae* to evaluate their response to four acaricides with different MoA: abamectin (GluCl allosteric modulator), bifenazate (METI-III), fenpyroximate (METI-I), and hexythiazox (MGI). The majority of the populations were collected from vegetables (eight out of twelve) while the remaining four populations (Tu1-Tu4) originated from citrus orchards. The results revealed strong variability in acaricide efficacy among populations (Table 2).

**Table 2.** Toxicity assays of four acaricides for ten *Tetranychus urticae* field populations from Greece.

| Acaricide class<br>(active ingredient) | GluCl allosteric modulators<br>(Abamectin) | METI-III<br>(Bifenazate) | METI-I<br>(Fenpyroximate) | MGI<br>(Hexythiazox) |
| --- | --- | --- | --- | --- |
| Population | Corrected Mortality at the recommended field dose (%Mortality ± SE) | | | |
| Tu1 | 98.47 ± 1.53 | 100.00 ± 0.00 | 92.22 ± 4.07 | 99.35 ± 0.65 |
| Tu2 | 100.00 ± 0.00 | 100.00 ± 0.00 | 95.89 ± 2.26 | 99.29 ± 0.71 |
| Tu3 | 100.00 ± 0.00 | 96.33 ± 1.89 | 87.97 ± 5.12 | 100.00 ± 0.00 |
| Tu4 | 98.51 ± 1.49 | 100.00 ± 0.00 | 98.63 ± 1.37 | 100.00 ± 0.00 |
| Tu5 | 36.99 ± 1.18 | 63.46 ± 9.67 | 9.46 ± 1.35 | 14.44 ± 2.09 |
| Tu6 | 100.00 ± 0.00 | 100.00 ± 0.00 | 100.00 ± 0.00 | 100.00 ± 0.00 |
| Tu9 | 24.6 ± 0.99 | 100.00 ± 0.00 | 28.83 ± 1.66 | 0.33 ± 0.58 |
| Tu10 | 13.10 ± 1.56 | 100.00 ± 0.00 | 19.66 ± 1.07 | 0.28 ± 1.38 |
| Tu11 | 100.00 ± 0.00 | 100.00 ± 0.00 | 100.00 ± 0.00 | 100.00 ± 0.00 |
| Tu12 | 100.00 ± 0.00 | 100.00 ± 0.00 | 100.00 ± 0.00 | 100.00 ± 0.00 |
| Tu13 | 100.00 ± 0.00 | 100.00 ± 0.00 | 100.00 ± 0.00 | 100.00 ± 0.00 |
| Tu14 | 100.00 ± 0.00 | 100.00 ± 0.00 | 100.00 ± 0.00 | 100.00 ± 0.00 |

Most tested populations (Tu1–Tu4, Tu6, Tu11–Tu14) exhibited high levels of susceptibility to all four compounds, with corrected mortality values ranging from 87.57% to 100%. However, three populations (Tu5, Tu9, and Tu10) showed reduced sensitivity to multiple acaricides. Tu5 showed especially low mortality to abamectin (36.99%), fenpyroximate (9.46%), and hexythiazox (14.44%) and low to moderate mortality to bifenazate (63.46%), suggesting resistance to several chemical groups. Similarly, Tu9 and Tu10 showed very low mortality to hexythiazox (0.33% and 0.28%), abamectin (24.6% and 13.10%) and fenpyroximate (28.83% and 19.66%), indicating multiple resistance traits.

Among the tested products, bifenazate was the most effective acaricide across the evaluated populations, consistently achieving 100% mortality in all except one (Tu5), where a moderate yet significant reduction in efficacy was observed (63.46%).

Five *T. urticae* populations were also tested against three alternative “green” products: Requiem, Eradicoat and Flipper (Table 3). Population Tu3 derived from citrus while the remaining four populations from vegetables. In general flipper and Requiem were the most effective products producing >70% mortality in four out of five populations. Population Tu10 showed high susceptibility to all three products, with mortality rates above 90%. Tu9 and Tu14 both showed high mortality to Requiem and Flipper (100%), but their response to Eradicoat was lower, with Tu9 showing reduced mortality (43.47%) and Tu14 showing a moderate response (71.93%). Tu3 showed moderate mortality to Requiem (74.64%) and Flipper (70.68%), but a low response to Eradicoat (23.58%). Tu5 was the least responsive, with very low mortality to Eradicoat (6.44%) and low response to Requiem (42.73%) and Flipper (39.27%).

**Table 3.** Toxicity assays of three green pesticides for five *Tetranychus urticae* field populations from Greece.

| Green pesticide | Requiem | Eradiocoat | Flipper |
| --- | --- | --- | --- |
| Population | Corrected Mortality at the recommended field dose (%Mortality ± SE) | | |
| Tu3 | 74.64 ± 2.58 | 23.58 ± 7.25 | 70.68 ± 4.67 |
| Tu5 | 42.73 ± 15.94 | 6.44 ± 5.41 | 39.27 ± 8.90 |
| Tu9 | 100.00 ± 0.00 | 43.47 ± 2.14 | 100.00 ± 0.00 |
| Tu10 | 91.97 ± 1.74 | 95.83 ± 1.62 | 100.00 ± 0.00 |
| Tu14 | 100.00 ± 0.00 | 71.93 ± 1.62 | 100.00 ± 0.00 |

### 3.2 Detection of resistance mutations to *T. urticae* field populations by ddPCR

Droplet digital PCR (ddPCR) analysis was used to quantify the frequency of resistance-associated target-site mutations in 17 *Tetranychus urticae* field populations. The results revealed substantial variability in resistance allele frequencies among populations and across different pesticide MoA (Table 4).

**Table 4.** Mutant allele frequencies measured by ddPCR in pools of field *T. urticae* populations.

| Acaricide class (active ingredient) | GluCl allosteric modulators (Abamectin) |  |  | Sodium channel modulators (Pyrethroids) |  | MGI (Etoxazole/Hexythiazox) | METI-I (Pyridaben) | METI-III (Bifenazate) |  |  |
| --- | --- | --- | --- | --- | --- | --- | --- | --- | --- | --- |
| Mutation | G314D | G326E | I321T | F1538 | L1024V | I1017F | H92R | G126S | S141F | P262T |
| Gene | GluCl1 | GluCl3 |  | VGSC |  | CHS1 | PSST | cytb |  |  |
| Population | %MAF (Mutant Allelic Frequency) |  |  |  |  |  |  |  |  |  |
| Tu1 | 0.00 | 0.00 | 0.00 | 0.00 | 0.92 | 1.89 | 0.00 | 0.00 | 0.00 | 0.00 |
| Tu2 | 0.00 | 0.00 | 0.39 | 0.00 | 0.74 | 0.40 | 0.00 | 0.41 | 0.00 | 0.00 |
| Tu3 | 0.00 | 0.00 | 0.00 | 0.00 | 0.00 | 0.00 | 0.00 | 0.00 | 0.00 | 0.00 |
| Tu4 | 0.00 | 0.00 | 0.00 | 0.00 | 0.00 | 0.00 | 0.00 | 0.00 | 0.00 | 0.00 |
| Tu5 | 0.00 | 0.00 | 54.45 | 100.00 | 0.00 | 100.00 | 0.74 | 100.00 | 0.00 | 0.00 |
| Tu7 | 0.00 | 0.00 | 30.04 | 0.00 | 0.00 | 0.00 | 0.00 | 0.00 | 0.00 | 0.00 |
| Tu8 | 0.00 | 0.00 | 16.10 | 1.65 | 0.00 | 0.00 | 0.00 | 0.00 | 0.00 | 0.00 |
| Tu9 | 0.00 | 0.00 | 100.00 | 100.00 | 0.00 | 100.00 | 0.00 | 100.00 | 0.00 | 0.00 |
| Tu10 | 0.00 | 0.00 | 100.00 | 0.00 | 0.00 | 100.00 | 0.00 | 100.00 | 0.00 | 0.00 |
| Tu12 | 0.00 | 0.00 | 0.00 | 100.00 | 0.00 | 0.00 | 0.00 | 0.00 | 0.00 | 0.00 |
| Tu13 | 0.00 | 0.00 | 0.00 | 4.64 | 0.00 | 0.00 | 0.00 | 0.00 | 0.00 | 0.00 |
| Tu14 | 0.00 | 0.00 | 0.00 | 0.00 | 0.00 | 0.00 | 0.00 | 0.00 | 0.00 | 0.00 |
| Tu15 | 0.00 | 0.00 | 0.00 | 31.41 | 0.00 | 0.00 | 0.00 | 0.00 | 0.00 | 0.00 |
| Tu16 | 0.00 | 0.00 | 69.04 | 92.33 | 0.00 | 91.17 | 0.00 | 92.79 | 0.00 | 0.00 |
| Tu17 | 0.00 | 0.00 | 0.00 | 0.00 | 0.00 | 0.00 | 0.00 | 0.00 | 0.00 | 0.00 |
| Tu18 | 0.00 | 0.00 | 10.07 | 0.00 | 0.00 | 0.00 | 0.00 | 0.00 | 0.00 | 0.00 |
| Tu19 | 0.00 | 0.00 | N/A | 42.71 | 0.00 | 100.00 | 47.77 | 100.00 | 0.00 | 42.89 |

Mutations associated with resistance to GluCl allosteric modulators (abamectin), including *G314D* and *G326E* in *GluCl1* and *GluCl3* respectively, were absent in all populations (0% MAF). The *I321T* mutation in *GluCl3*, also linked to abamectin resistance, was detected in eight populations, in fixation (100%) in two populations (Tu9, Tu10), in high frequencies in Tu5 (54.45%), Tu16 (69.04%), and lower levels in Tu7 (30.04%), Tu8 (16.10%), Tu18 (10.07%), and Tu2 (0.39%).

For sodium channel modulators (pyrethroids), *F1538I* was widely present (8 populations), being fixed or nearly fixed in Tu5 (100%), Tu9 (100%), Tu12 (100%), and Tu16 (92.33%). Intermediate or lower frequencies were observed in Tu15 (31.41%), Tu19 (42.71%), Tu13 (4.64%), and Tu8 (1.65%). In contrary, *L1024V* was only detected at extremely low frequency in Tu1 (0.92%) and Tu2 (0.74%) both originating from citrus.

The *I1017F* mutation in *CHS1*, conferring resistance to mite growth inhibitors (MGIs) such as etoxazole and hexythiazox, was highly prevalent or fixed in five populations (Tu5, Tu9, Tu10, Tu16 and Tu19), and present at very low levels in Tu1 (1.89%) and Tu2 (0.40%).

Regarding mitochondrial electron transport inhibitors (METIs), *H92R* (associated with METI-I resistance) was detected at moderate frequency in populations Tu19 (47.77%) and at extremely low frequency in Tu5 (0.74%). *G126S* was highly prevalent or fixed in Tu5, Tu9, Tu10, Tu16, and Tu19 while found at low levels in Tu2 (0.41%). The *P262T* mutation was detected only in Tu19 (42.89%), while *S141F* was not detected in any population.

No resistance mutations were detected in four populations (Tu3, Tu4, Tu14 and Tu17) suggesting that these populations are fully susceptible based on the tested markers. On the other hand, Tu5, Tu9, Tu10, Tu16, and Tu19 harbored multiple high-frequency resistance mutations, indicating multi-resistant phenotypes.

### 3.3 Detection of resistance mutations to *B. tabaci* field populations by ddPCR

Six *Bemisia tabaci* field populations (Bt1–Bt6) were screened using droplet digital PCR (ddPCR) for key target-site resistance mutations associated with multiple insecticide classes (Table 5).

**Table 5.** Mutant allele frequencies measured by ddPCR in pools of B. tabaci field populations.

| Acaricide class (active ingredient) | AChE inhibitors (OPs, carbamates) | Sodium channel modulators (Pyrethroids) |  | Lipid biosynthesis inhibitors targeting ACC (tetronic & tetramic acid derivatives / ketoenols)) |
| --- | --- | --- | --- | --- |
| Mutation | F331W | L925I | T929V | A2083V |
| Gene | *Ace1* | *VGSC* |  | *Acc* |
| Population | %MAF (Mutant Allelic Frequency) |  |  |  |
| Bt1 | 92.44 | 34.36 | 58.24 | 88.41 |
| Bt2 | 89.64 | 35.72 | 56.48 | 89.82 |
| Bt3 | 21.48 | 2.90 | 10.73 | 1.39 |
| Bt4 | 95.56 | 31.65 | 51.83 | 71.43 |
| Bt5 | 89.30 | 37.49 | 57.94 | 78.12 |
| Bt6 | 92.11 | 44.74 | 45.29 | 92.74 |

The *F331W* mutation in the *Ace1* gene, which confers resistance to acetylcholinesterase inhibitors (organophosphates and carbamates), was detected at high frequencies in all populations except Bt3 (21.48%), ranging from 89.30% in Bt5 to 95.56% in Bt4. For pyrethroid resistance, both *L925I* and *T929V* mutations in the *vgsc* gene were consistently detected. The combined MAFs for these two mutations ranged low levels in Bt3 (*L925I*: 2.90%, *T929V*: 10.73%) to high levels in Bt6 (*L925I*: 44.74%, *T929V*: 45.29%). The *A2083V* mutation in the *acc* gene, associated with resistance to lipid biosynthesis inhibitors (tetronic and tetramic acid derivatives), showed strong variation. It was found to have a very high frequency in five out of six populations (Bt1, Bt2, Bt4, Bt4 and Bt5) ranging from 71.43% to 92.74% while detected at extremely low frequency in population Bt3 (1.39%).

Among the tested populations, Bt6 appeared as the most resistant, showing the highest MAFs across all tested classes: *F331W* (92.11%), *L925I*/*T929V* (44.74/45.29%), and *A2083V* (92.74%), indicating multiple target-site resistance. In contrast, Bt3 exhibited the lowest MAFs, *F331W* (21.48%), *L925I*/*T929V* (2.90/10.73%), and *A2083V* (1.39%), suggesting a more susceptible profile.

## 4. Discussion

This study focused on field populations of *Tetranychus urticae* and *Bemisia tabaci* collected from major agricultural regions across Greece, including both mainland areas and the island of Crete. These regions are characterized by intensive fruit and vegetable production, in both open-field and greenhouse conditions, where pest control relies heavily on repeated chemical applications. Such practices exert strong selection pressure on pest populations, facilitating the development and spread of resistance.

The toxicity data for four acaricides (abamectin, bifenazate, fenpyroximate, and hexythiazox) demonstrated a wide range of responses among *T. urticae* populations. Among the twelve populations tested, four originated from citrus orchards and eight from vegetable crops. Citrus populations were almost fully susceptible to all tested acaricides, whereas populations from vegetables showed varying responses at the recommended field dose. Although the number of populations tested is limited and the application history is incomplete, this variability may be associated with the host plant and/or treatment practices. Three populations from vegetables, collected in Messinia (Peloponnese) and Tympaki (Crete), displayed low sensitivity to three or all four tested acaricides, indicating multi-resistant phenotypes. The remaining five populations from vegetables were fully susceptible to all tested compounds, consistent with the known pesticide use history in the field. In our previous study (Mavridis et al., 2021), more than half of the tested populations were found to be multi-resistant to the majority of the acaricides. This difference may be attributed to the fact that those populations were collected from Crete five years ago, from heavily treated greenhouse systems growing vegetables and ornamentals.

Additionally, the toxicity of three “green” products (Requiem, Eradicoat, and Flipper) was evaluated in five populations with different host plants and varying responses to synthetic pesticides. One population was collected from citrus (Tu3), which was susceptible to all acaricides, while four were collected from vegetables and included both multi-resistant (Tu5, Tu9, Tu14) and susceptible (Tu10) populations.

A panel of twelve target-site resistance mutations, previously developed and applied in (Mavridis et al., 2022b), was used to detect resistance in *T. urticae*. These assays allowed us to identify multiple mutations associated with resistance, some of which have also been linked to phenotypic resistance and provided useful comparisons with previous findings.

Mutations in GluCl1 (G314D) and GluCl3 (G326E and I321T) have been strongly associated with abamectin resistance (Dermauw et al., 2012; Papapostolou et al., 2021; Xue et al., 2020). Among the three, only I321T was detected in our study, in eight populations with varying frequencies, whereas G314D and G326E were completely absent. In populations Tu5, Tu9, and Tu10 — where both toxicity and molecular data were available — there was a clear correlation between low mortality and high mutation frequency. Compared to our previous study (Mavridis et al., 2022b), where G326E alone or in combination with G314D was detected in four out of six populations and associated with low mortality at RD (as low as 6%), the present data show a shift toward the prevalence of I321T.

F1538I and L1024V are known pyrethroid resistance mutations (Kwon et al., 2010; A. Tsagkarakou et al., 2009). L1024V was detected at very low frequency in only two citrus populations (<1%), while F1538I was more frequent, found in nearly half of the tested populations and at high frequency in several cases. This prevalence is consistent with our previous findings (Mavridis et al., 2022b). Although pyrethroids are not registered against *T. urticae* in citrus or vegetables, their use against other insect pests may explain the selection pressure and the observed frequencies.

The I1017F mutation in CHS1 was found at high frequency in populations with low hexythiazox mortality, confirming its strong link to MGI resistance. It was found fixed in Tu5, Tu9, and Tu10, all of which showed poor response to hexythiazox (mortality at RD below 15%). These results agree with (Mavridis et al., 2022b), where this mutation was detected at high frequency in half of the populations and was also correlated with resistance to etoxazole.

H92R, a mutation in PSST associated with resistance to METI-I acaricides (e.g., pyridaben, fenpyroximate), was detected in only two populations — one at moderate (48%) and one at very low frequency (<1%). Interestingly, toxicity data showed low mortality to fenpyroximate in three populations (Tu5, Tu9, Tu10), although the H92R mutation was not present. In our previous study (Mavridis et al., 2022b), H92R was frequently found in Crete populations, though it showed poor association with pyridaben toxicity.

For bifenazate resistance, three cytb mutations (G126S, S141F, and P262T) were examined. G126S was found at high or fixed frequency in several populations, while P262T was detected in only one case, in combination with G126S. S141F was not detected in any sample. In our previous study, only G126S was detected, consistent with the current findings — apart from Tu19, where P262T was also identified. To our knowledge, this is the first detection of P262T in Greece and may reflect evolving resistance to bifenazate. Previous research has shown that G126S alone does not confer resistance (Xue et al., 2021).

Overall, the consistency between molecular and bioassay data supports the value of these markers, while the differences from earlier studies indicate ongoing resistance evolution.

In *B. tabaci*, most populations carried multiple target-site mutations: F331W (ace1), L925I/T929V (vgsc) and A2083V (acc). The F331W mutation, which was previously fixed in Crete populations (A. Tsagkarakou et al., 2009), was still present at high frequency (>89%) in all samples except Bt3 (21.48%). The reduction in frequency could reflect the reduced use of organophosphates and the high fitness cost of this mutation.

Similarly, both L925I and T929V mutations were found at moderate to high frequencies in all populations except Bt3. In our previous survey (Mavridis et al., 2022a), L925I was more frequent than T929V, whereas here T929V appears slightly more prevalent — possibly due to their mutually exclusive nature. Earlier studies in Greece and other Mediterranean countries (Gauthier et al., 2014; Anastasia Tsagkarakou et al., 2009) have also reported high frequencies of kdr mutations, likely due to decades of pyrethroid use and the absence of major fitness costs.

The A2083V mutation, associated with ketoenol resistance, was found at high to very high frequencies (>71%) in five out of six populations, and at very low frequency in Bt3. In our previous work, this mutation was detected in 8 of 11 populations, but usually at lower levels (>70% in only 3 populations). Although we did not perform spiromesifen bioassays in the current study, our earlier findings (Mavridis et al., 2022a) (Mavridis et al., 2022) support a strong genotype–phenotype link for A2083V.

Although this study focused on target-site resistance mechanisms, metabolic resistance — such as overexpression of detoxification enzymes — is also known to play an important role in both species and should be considered in future surveys.

In conclusion, our study confirms the presence of widespread resistance in *Tetranychus urticae* and *Bemisia tabaci* populations in Greece, with strong associations between bioassay results and key resistance mutations. The detection of multiple target-site mutations in the same populations suggests increasing multi-resistance and reduced pesticide efficacy. These findings highlight the need for regular molecular resistance monitoring, rotation of pesticide modes of action, and greater integration of alternative tools such as biopesticides, to support effective and sustainable IPM strategies.

